# Genome-Wide Fetalization of Enhancer Architecture in Heart Disease

**DOI:** 10.1101/591362

**Authors:** Cailyn H. Spurrell, Iros Barozzi, Brandon J. Mannion, Matthew J. Blow, Yoko Fukuda-Yuzawa, Sarah Y. Afzal, Jennifer A. Akiyama, Veena Afzal, Stella Tran, Ingrid Plajzer-Frick, Catherine S. Novak, Momoe Kato, Elizabeth Lee, Tyler H. Garvin, Quan T. Pham, Anne N. Harrington, Steven Lisgo, James Bristow, Thomas P. Cappola, Michael P. Morley, Kenneth B. Margulies, Len A. Pennacchio, Diane E. Dickel, Axel Visel

## Abstract

Heart disease is associated with re-expression of key transcription factors normally active only during prenatal development of the heart. However, the impact of this reactivation on the genome-wide regulatory landscape in heart disease has remained obscure. Here we show that pervasive epigenomic changes occur in heart disease, with thousands of regulatory sequences reacquiring fetal-like chromatin signatures. We used RNA-seq and ChIP-seq targeting a histone modification associated with active transcriptional enhancers to generate genome-wide enhancer maps from left ventricle tissue from 18 healthy controls and 18 individuals with idiopathic dilated cardiomyopathy (DCM). Healthy individuals had a highly reproducible epigenomic landscape, consisting of more than 31,000 predicted heart enhancers. In contrast, we observed reproducible disease-associated gains or losses of activity at more than 7,500 predicted heart enhancers. Next, we profiled human fetal heart tissue by ChIP-seq and RNA-seq. Comparison with adult tissues revealed that the heart disease epigenome and transcriptome both shift toward a fetal-like state, with 3,400 individual enhancers sharing fetal regulatory properties. Our results demonstrate widespread epigenomic changes in DCM, and we provide a comprehensive data resource (http://heart.lbl.gov) for the mechanistic exploration of heart disease etiology.

## Introduction

Heart failure resulting from cardiomyopathy is a prevailing cause of adult mortality worldwide^1^. Among cardiomy-opathies-dilated cardiomyopathy (DCM) is the most common^2^ and is characterized by the dilation and weakening of the left ventricle and decreased ejection fraction in the absence of coronary artery disease or other abnormalities^3^. Approximately one third of DCM cases are explained by coding mutations in known genes^4,5^. Gene expression studies have been widely used to facilitate the identification of additional causal genetic variation and to understand the transcriptional pathways impacted during heart failure, which may provide new therapeutic targets^6–8^. Transcriptional profiling of failing hearts, including those with idiopathic DCM, has identified numerous coding and non-coding genes whose expression is altered in heart disease^7,9–12^. These studies have consistently shown that heart failure is associated with the reactivation of a fetal gene program: a curated set of genes with well-established roles in fetal heart development that are normally repressed in non-failing adult heart^13,14^. Notably, a fetalization mechanism has also been suggested for similar pathways involved in lung disease^15,16^ and tumorigenesis^17^. However, the full quantitative extent of fetal gene program reactivation in disease processes and the identity of the gene regulatory sequences correlated with these changes remain unknown.

Gene expression is controlled in part by enhancers, a major class of cis-regulatory elements, which can act over large genomic distances to activate gene transcription in specific cell types or developmental stages ^9^. Chromatin immuno, precipitation followed by sequencing (ChIP-seq) targeting enhancer-associated chromatin marks (e.g., H3K27ac) can be used to generate accurate genome-wide maps of enhancers active in a tissue of interest^10,18^. However, the small number of ChIP-seq data sets from individual human heart samples described to date has been insufficient to systematically assess the reproducibility of the cardiac enhancer landscape in healthy individuals or to quantify and characterize possible changes associated with disease states^19–21^. In particular, a recent study examined cardiomyocyte cells isolated from a small number of failing and non-failing hearts to explore the regulatory landscape of these cells in disease^22^ but identified a very limited set of enhancers that were differentially active between these states.

We performed a comprehensive exploration of human heart enhancers in left ventricle samples from an extensive cohort of control adults and individuals with dilated cardiomyopathy (see study overview in **Figure 1**). In healthy hearts, we observe a highly reproducible enhancer landscape across individuals. Furthermore, we observe major changes in the regulatory landscape of DCM samples and, through comparison with fetal heart tissue samples, demonstrate that more than 3,400 enhancers adopt fetal-like regulatory activity states in DCM. We also show that these enhancers are enriched in a specific subset of basic helix-loop-helix (bHLH) transcription factor binding sites and are sufficient to drive cardiac expression *in vivo* in transgenic mouse reporter assays. We demonstrate, through correlation with transcriptome data, that enhancers activated in disease are in close proximity to genes with increased expression in disease. Finally, we identify more than 200 fetal heart genes that are up-regulated in adult heart failure. Taken together, our findings indicate that a large number of fetal heart enhancers become consistently reactivated in heart disease, and our results identify the genomic locations and activation states of these enhancers, providing a rich resource for the exploration of mechanistic links between enhancers and clinical cardiac phenotypes.

**Fig. 1.**
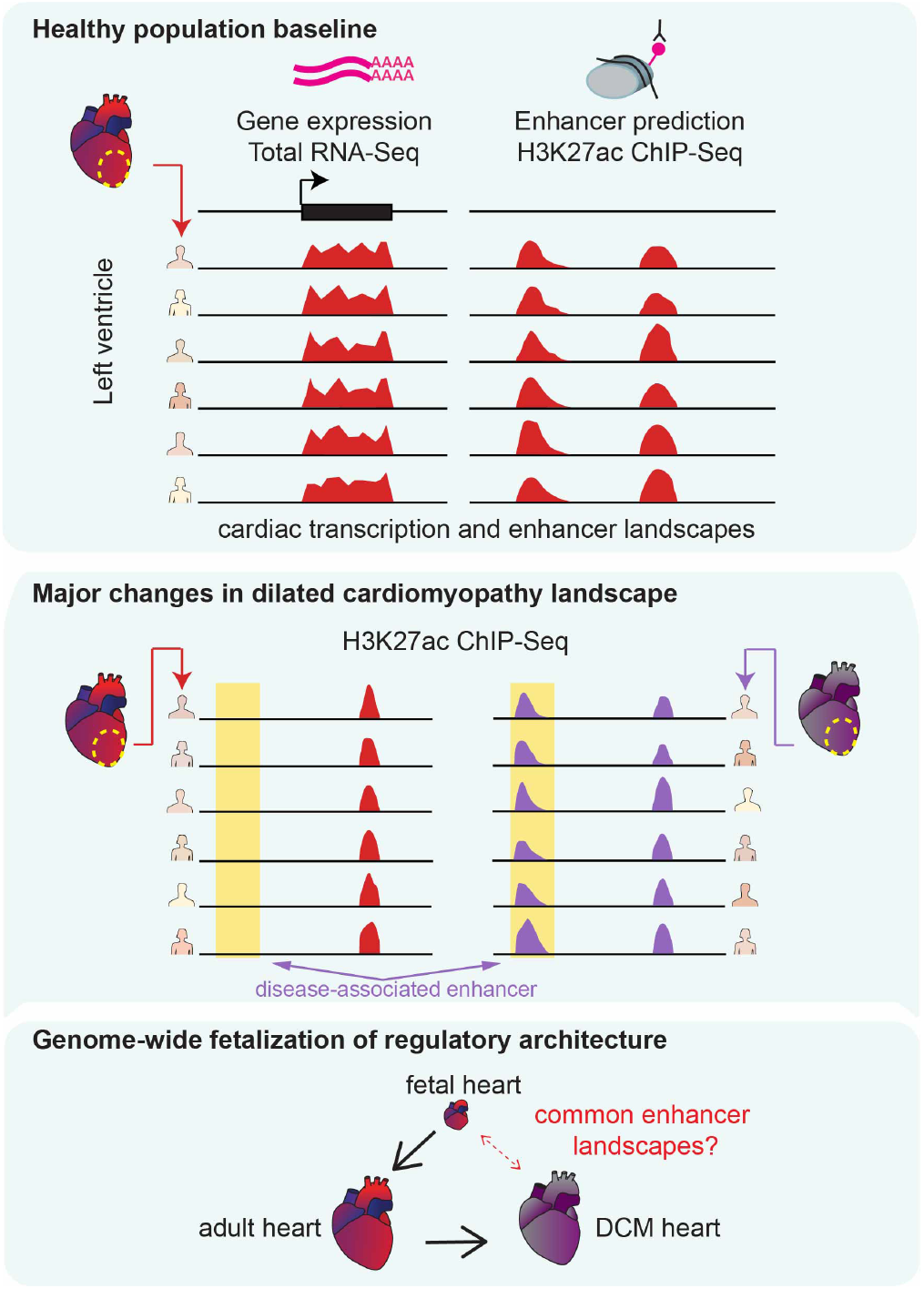
Overview of this study. We performed RNA-seq and ChIP-seq on left ventricle samples from 18 healthy donors. We identify reproducible enhancer predictions and gene expression patterns in healthy heart (top). Next, we identify global alterations in enhancer occupancy from 18 hearts with idiopathic dilated cardiomyopathy (DCM) (center). Last, we compare adult healthy and disease states with 5 embryonic or fetal heart and identify similarities in the enhancer landscapes of DCM and fetal heart (bottom).

## Results

### Highly Reproducible Heart Enhancer Landscape in Healthy Individuals

To assess the heterogeneity of the gene regulatory landscape in healthy human hearts, we used chromatin profiling to identify putative enhancer regions. We performed ChIP-seq with an antibody to H3K27ac on left ventricle samples from 10 male and 8 female healthy adult donor hearts for which an acceptable transplant recipient could not be found (**Figure 1, Supplementary Table 1**). In parallel, we performed RNA-seq on these samples to enable a comparison of each enhancer profile with its associated transcriptome. We defined enhancers as genomic regions with significant H3K27ac enrichment that are at least 1 kbp away from the nearest transcription start site (see **Methods**). All chromatin and RNA-seq data from these samples, as well as disease samples described below, are available for download and browsing in an accompanying web resource (heart.lbl.gov). Control samples had an average of 17,352 putative distal enhancers, with 40% of peaks identified in any given control sample present in all other control samples, 78% shared across two-thirds of all control samples, and only 9% (685) unique to the individual (**Figure 2a**). Across all 18 control samples, we identified a total of 43,363 predicted distal enhancers present in at least one sample, with more than 70% (31,033) found in at least two individuals (**Supplementary Figure 1, Supplementary Table 2**). To determine the functions of genes associated with the cardiac enhancers predicted through this approach, we examined the enrichment of functional ontology terms ^24^ among genes located near candidate regions showing strong H3K27ac enrichment in at least 16 healthy controls (see **Methods**). Among the most significantly enriched annotations derived from mouse gene expression studies, 11 of the top 20 are cardiovascular-related functions (**Supplementary Table 3**). Our results indicate that the enhancer landscape in healthy human hearts is overall highly reproducible, with quantitatively modest differences between individuals.

**Fig. 2.**
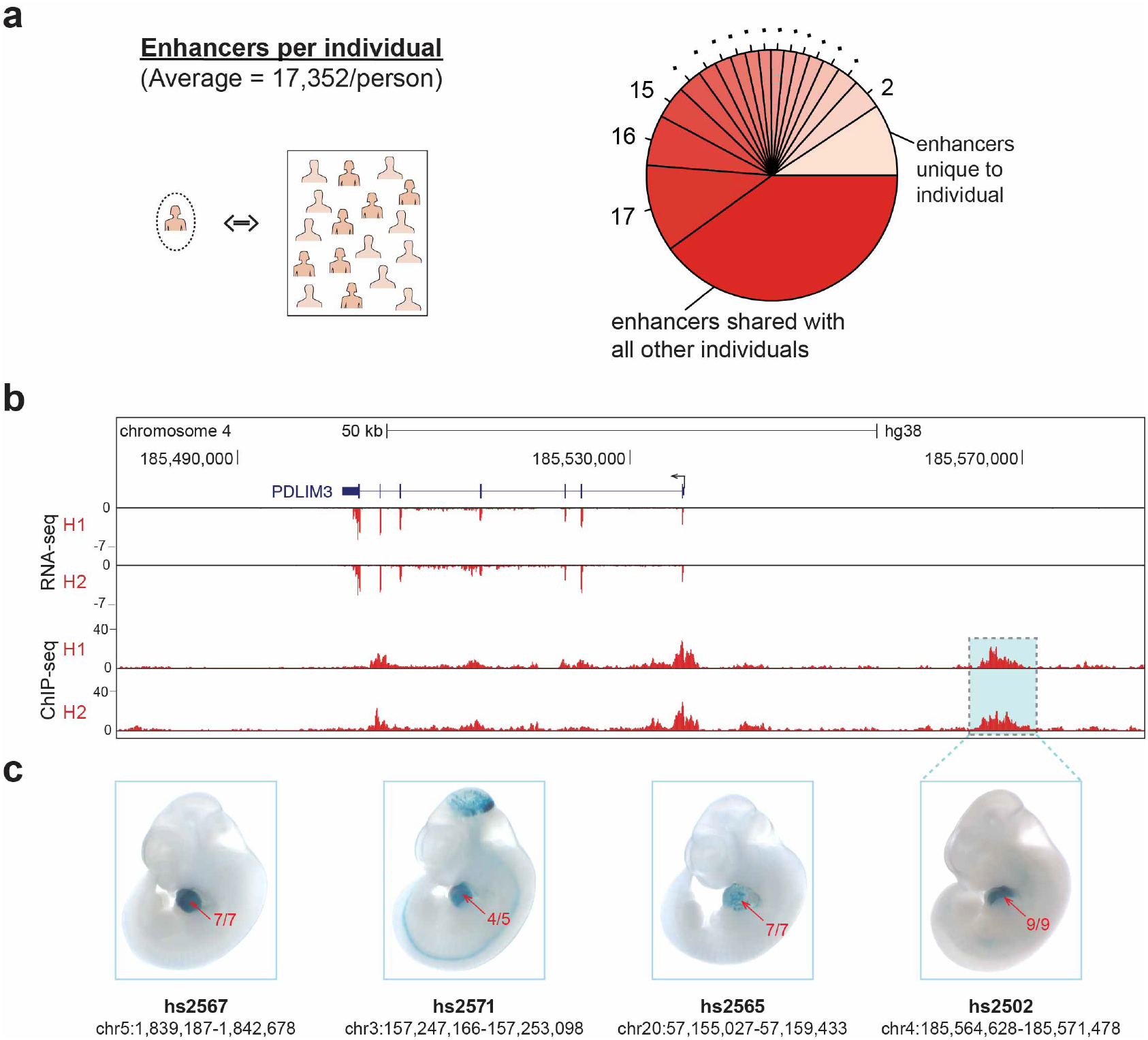
The cardiac enhancer landscape is highly reproducible across individuals. **a,** Average proportion of peaks from a given control sample shared with other control samples. **b,** Paired RNA-seq and ChIP-seq tracks from two samples, H1 and H2, at the *PDLIM3* locus. **c,** Transgenic mouse assay validation of four heart enhancers, including one upstream of *PDLIM3* (see Supplementary Figure 2 for results for additional predicted heart enhancers). One representative embryonic day 11.5 embryo is shown for each enhancer, and numbers in red show the reproducibility of heart staining in each transgenic experiment. Information underneath each embryo includes the identification number in the VISTA Enhancer Browser^23^ (enhancer.lbl.gov) and the human genome coordinates (hg38) of each tested enhancer.

To verify that enhancers predicted by this ChIP-seq approach are bona fide cardiac-specific regulatory sequences, we first compared our predicted human heart enhancers with the VISTA Enhancer Browser (enhancer.lbl.gov), an existing catalog of *in vivo* validated enhancers^23^. This database contains 283 human and mouse loci that have been tested using transgenic mouse assays and have validated activity in the heart. Nearly 55% (n=154) of these validated heart enhancers overlapped peaks in our adult human heart samples (**Supplementary Table 2**). We additionally performed forward *in vivo* validation using transgenic mouse reporter assays^25^ at embryonic day 11.5 for several candidate human adult heart enhancers. Examples of reproducible in vivo cardiac reporter activity driven by different candidate enhancers, including one near *PDLIM3*, are shown in **Figure 2b, c.** Together, these results reinforce that predicted adult heart enhancers drive gene expression in heart tissue.

### Genome-wide Changes in Enhancer Architecture in DCM

We next evaluated possible global changes in enhancer architecture in heart disease using transcriptome and chromatin profiling on explanted left ventricle samples from individuals with late stage idiopathic DCM. ChIP-seq data were generated from 18 DCM cases that were age- and sex-matched to the 18 healthy control samples (see **Methods, Supplementary Table 1**), and we obtained RNA-seq data from 15 of the 18 DCM left ventricle samples. Across the two sets of samples (healthy and DCM), the greatest driver of variance is disease state, explaining 18.1% of the variability in gene expression and 18.3% of the variability in predicted enhancers (**Figure 3a, b**). Biological sex was not identified as a substantial driver of variance for either control or DCM samples (**Supplementary Figure 3**). Of the 18,913 genes expressed in at least two individuals, 870 (4.6%) were differentially expressed between disease and health, with 416 up-regulated in DCM and 454 downregulated in DCM (P <0.01, |log_2_(fold change)| ≥ 1; **Figure 3c, Supplementary Table 4**). From the combined pool of 36 left ventricle H3K27ac ChIP-seq samples, we identified 38,883 predicted enhancers present in at least two individuals (**Supplementary Table 5**). Our large sample size provided sufficient power to use differential peak analysis^26^ to identify enhancers with quantitative differences between healthy and DCM samples that could be contributing to or resulting from the observed gene expression differences. We identified 7,507 (19%) enhancers that are differentially bound between healthy and DCM samples (P < 0.01 and |log_2_(fold change)| ≥ 1; DiffBind DESeq analysis, see **Methods** and **Supplementary Table 6**). Among these differential peaks, 4,406 show increased signal in DCM relative to healthy controls, whereas 3,101 have increased predicted enhancer activity in controls (**Figure 3c, Supplementary Table 6**). These results show that, in addition to the dysregulation of hundreds of cardiac-expressed genes, the genome-wide enhancer architecture is substantially altered in left ventricle from DCM patients.

**Fig. 3.**
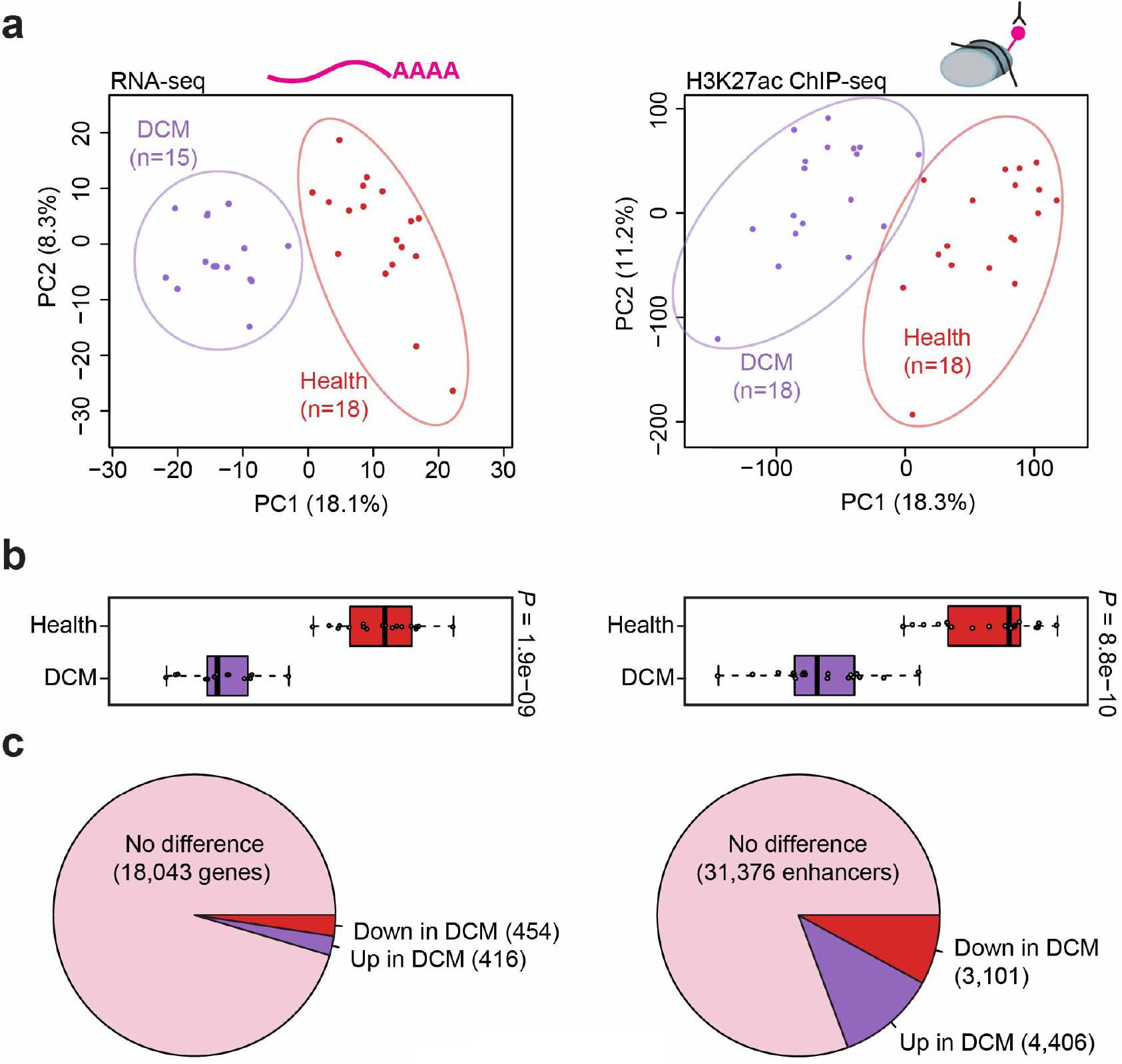
Extensive reprogramming of the epigenomic architecture in heart disease. **a,** Principal component analysis showing the first two principal components (PC1 and PC2) for the top 1000 variably expressed genes from RNA-seq (left) and for distal enhancer peaks from ChIP-seq (right). Each point indicates a unique sample, color coded by cohort. **b,** Boxplots showing PC1 for each sample by cohort (left: RNA-seq, right: ChIP-seq). Boxplots indicate median and quartile values for each data set; points indicate individual samples. PC1 separates disease state in both RNA-seq and ChIP-seq data (P values by two-sided Mann-Whitney U test). **c,** Pie charts showing the proportion of unchanged versus differentially expressed genes (left) and differentially bound enhancer peaks (right) relative to each cohort (see **Methods** for details).

### Differential Regulation of Disease-Related Genes and Pathways

To further assess the regulatory relationship between differentially active enhancers and differentially expressed genes in DCM, we examined correlations between RNA-seq and ChIP-seq data. We observed multiple anecdotal examples of differentially active enhancers located near genes with the same directionality of differential expression. An illustrative example is provided by the *SMOC2* locus, where a strong, DCM-specific increase in H3K27ac signal at a site 20 kb upstream of the promoter correlates with a strong increase in *SMOC2* expression in DCM (**Figure 4a**). To examine this effect beyond individual loci, we performed a global correlation analysis, assessing the enrichment of differentially bound candidate enhancers near differentially expressed genes. Differentially bound candidate enhancers with increased binding are indeed enriched near differentially upregulated genes, and candidate enhancers with decreased binding are enriched near downregulated genes, with the most pronounced enrichment observed within the 100 kb window closest to the promoter (**Figure 4b, c**).

**Fig. 4.**
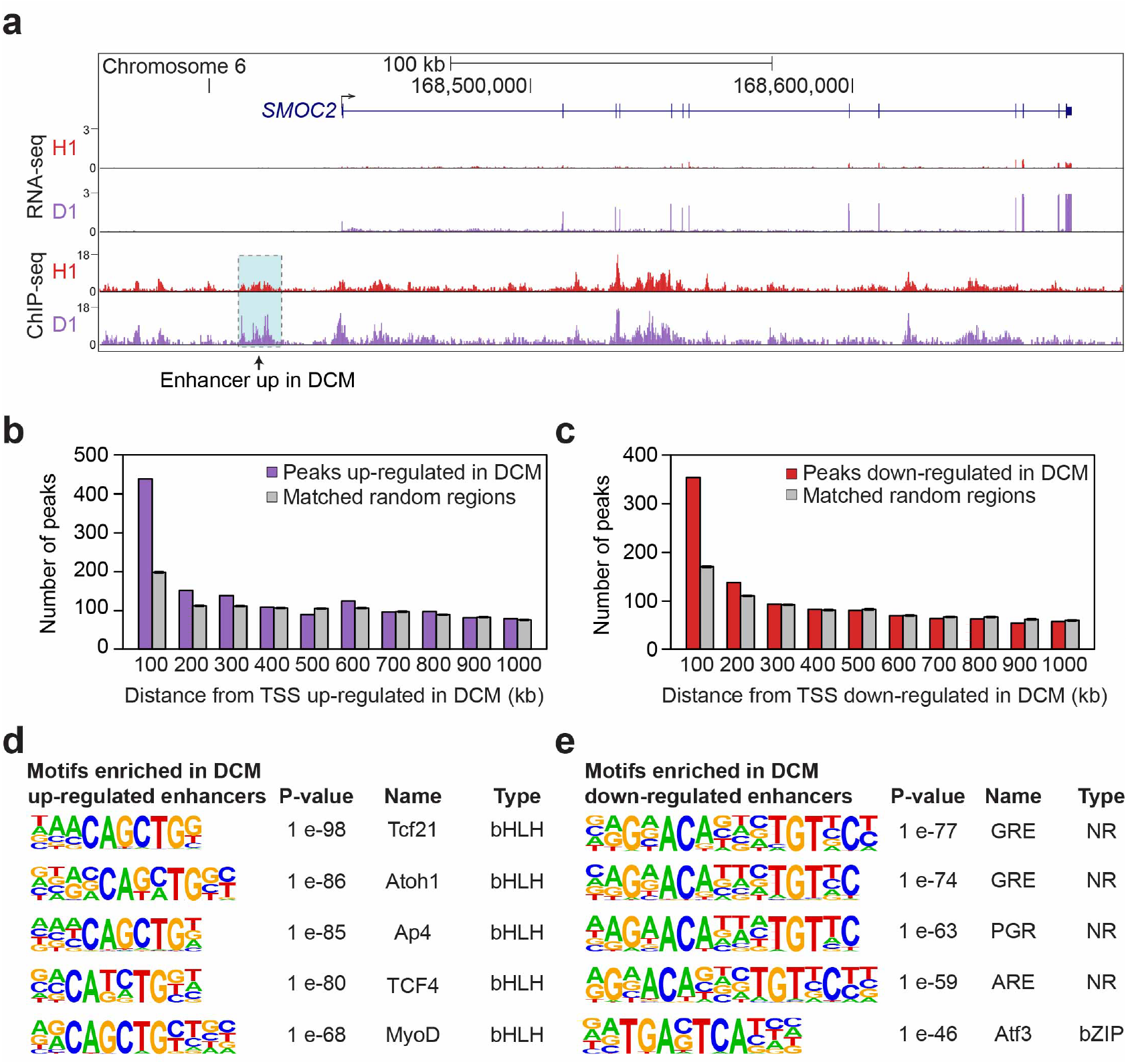
Differentially bound distal enhancers associated with heart failure-specific gene expression changes. **a,** Co-occurrence of increased expression of *SMOC2* (by RNA-seq) and increased H3K27ac at a nearby enhancer (by ChIP-seq) in DCM. Shown are data from a representative healthy (H1) and DCM (D1) sample. **b,** Distal ChIP-seq peaks relative to their distance from transcription start sites (TSS) of genes up-regulated in DCM, divided into 100 kb bins. Purple: distal peaks up-regulated in DCM. Gray: a random subset of 4,406 regions matched to all heart ChIP-seq peaks was assessed. The average across 200 sets of randomized control elements is shown. **c,** Same analysis as in **b,** but for genes with decreased expression in DCM and distal peaks down-regulated in DCM (red). A random subset of 3,101 regions matched to all heart ChIP-seq peaks was assessed (gray). **d,e,** Transcription factor binding sites enriched in peaks up-regulated (**d**) and down-regulated (**e**) in DCM^28^. P values by HOMER.

To assess the biological functions of the differentially expressed genes and differentially active enhancers, we examined their enrichment for biological process ontology terms^24^. For the 4,406 enhancer peaks with predicted increased activity in DCM, there is enrichment for terms including extracellular matrix remodeling (**Supplementary Table 7**). Similarly, genes up-regulated in DCM independently show enrichment for extracellular matrix components and cell adhesion (**Supplementary Table 8**). These results reinforce our conclusion that DCM-specific enhancers are enriched near genes that are differentially regulated in disease, highlighting fibrosis pathways in heart failure^27^.

To identify candidate transcription factors (TFs) that may be driving altered enhancer activity in DCM, we performed binding site enrichment analysis^28^ for distal enhancer peaks with increased or decreased activity in DCM. In peaks with increased activity in DCM, binding sites for bHLH transcription factors are enriched (**Figure 4d**). Several of the most significant factors in this class, including TCF4 and TCF21, are involved in epithelial-to-mesenchymal transition^29^, which may contribute to cardiac fibrosis^30^. In peaks with decreased activity in DCM, there is enrichment in nuclear receptor transcription factor binding sites (**Figure 4e**). We found particularly pronounced enrichment in glucocorticoid response elements (GRE), which are involved in growth and immune response pathways^31^. We also found enrichment of bZIP factors that are part of the AP-1 complex, which has been suggested to act as homeostatic regulator required to maintain a steady state of cell proliferation^32,33^. Taken together, these data show that heart disease-associated enhancers are enriched near genes differentially regulated in disease and that known functions of these genes are consistent with roles in heart failure.

### Fetalization of the Regulatory Landscape in Heart Disease

Previous studies have reported that some genes that are essential for fetal heart development are downregulated in adult heart, but become reactivated in failing hearts^7,13,34^. However, beyond anecdotal examples, the transcriptome-wide scale of such re-expression of prenatally expressed human genes in DCM has remained unclear. Likewise, it is unknown whether reactivation of these fetal genes is associated with reactivation of fetal regulatory elements. To comprehensively define the fetal gene program reactivated in dilated cardiomyopathy and to test whether enhancer regulation correlates with the re-expression of fetal genes, we performed ChIP-seq and RNA-seq on five embryonic and fetal human heart samples, covering developmental stages ranging from 8 weeks to 17 weeks post conception (**Figure 5a, Supplementary Tables 9,10**; see **Methods** for sample details). We performed principal component analysis followed by dimensionality reduction using t-SNE to capture variation from multiple principal components for healthy, DCM, and prenatal samples (**Figure 5b, c; Supplementary Figure 4**; and **Methods**). These analyses, capturing the local and global structure of complex data for visualization in two dimensions ^35^, indicate that the three groups of samples (adult controls, adult DCM, fetal) show distinct genome-wide transcriptome and enhancer signatures (**Figure 5b, c**). Consistent with the notion of reactivation of fetal genes in heart disease, we found that genes expressed during fetal development are enriched among those upregulated in DCM relative to healthy adult heart and vice versa (**Supplementary Figure 4c, d**).

**Fig. 5.**
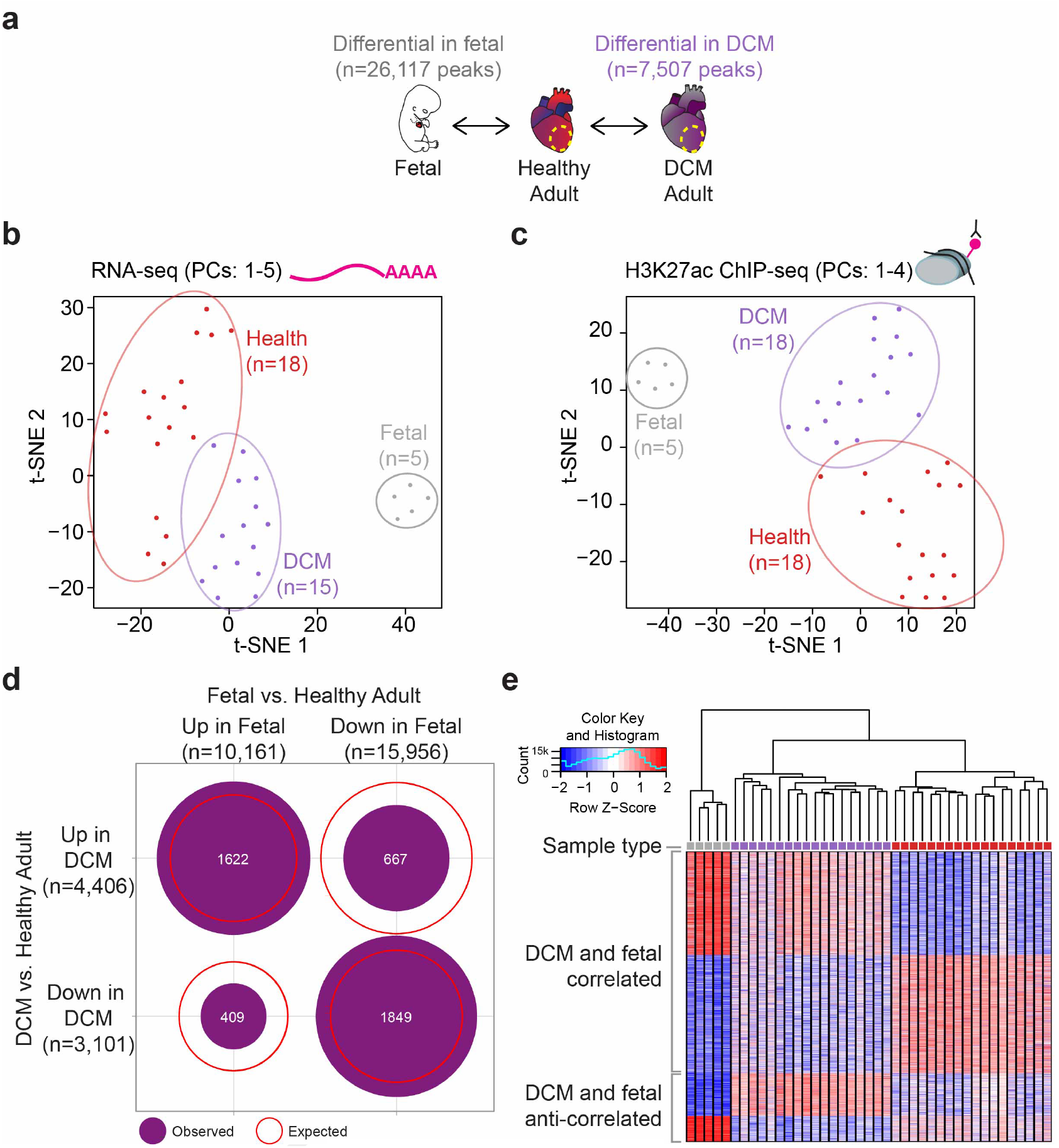
Fetalization is a major driver of regulatory changes in heart disease. **a,** Schematic of experimental comparisons: healthy adult peaks vs. healthy fetal peaks, and healthy adult peaks vs. DCM adult peaks. **b,** Dimensionality reduction visualization of gene expression shown by t-SNE. The t-SNE incorporates data from the top principal components (selected by jackstraw cutoff, perplexity=10). Prenatal samples are gray, DCM samples are purple, and healthy samples are red. **c,** Same analysis as **b,** for H3K27ac-predicted enhancers. **d,** Number of enhancer peaks co-regulated in prenatal and DCM states. Purple circles show number of peaks overlapping in each quadrant. Red lines show expected overlap between each category. **e,** Heat map showing all differential peaks between 41 samples: 5 prenatal (gray), 18 DCM (purple), and 18 healthy (red).

In total, of the 416 genes upregulated in DCM, 213 are up-regulated in fetal heart relative to healthy adult. To examine if this coincides with enhancers returning to a fetal-like state in heart disease, we focused on the activity states of 4,547 regions that showed differential H3K27ac binding in both health versus DCM, as well as in adult versus fetal samples. The majority of these enhancers (76%, n=3,471) show the same activity state in fetal and adult DCM heart, respectively, resulting in an overlap of DCM- and fetal-biased peaks that was enriched 1.6-fold relative to the overlap expected by chance (**Figure 5d, e**). For the 1,622 enhancers with predicted increased activity in DCM and fetal, there is enrichment for gene ontology terms associated with fibrosis and extracellular matrix remodeling, along with terms involving TGF*β* signaling (**Supplementary Table 11**). Genes in these pathways are upregulated in cardiac development and in heart failure^36,37^.

In addition to enhancers that showed fetal-like regulatory properties in DCM, we also observed populations of enhancers that are either anti-correlated with fetal enhancer activity states (n=1,076) or that were DCM-specific, i.e. showed activity states that were different from fetal and healthy adult cardiac tissue (n=2,960). Both of these groups of enhancers indicate that while the genome-wide alterations in enhancer landscape in DCM are dominated by a shift toward fetal-like signatures, there are clearly additional, DCM-specific molecular events, as expected from the pathological context associated with heart disease. To provide additional insight into gene regulatory processes that are shared between DCM and fetal development versus those that are specific to DCM, we examined enrichment of transcription factor binding site motifs in the 1,622 enhancers that have increased activity in both versus the 2,784 that are upregulated only in DCM. Both sets of enhancers were generally enriched for bHLH transcription factor binding motifs, but we observed the most significant enrichment for distinct subsets of bHLH motifs (**Supplementary Figure 5**). For example, enhancers with increased activity in both DCM and fetal samples were especially enriched in motifs for MyoD and Myf5, proteins with well-established roles in myogenesis^38^. Taken together, our results support that the upregulation of fetal genes in DCM is accomplished through the reactivation of fetal enhancers. Importantly, the genome-wide expression and enhancer maps generated through this study will enable the targeted follow-up of regulatory events associated with the pervasive reactivation of fetal expression programs, as well as molecular downstream events specifically associated with disease progression.

## Discussion

Understanding the mechanisms regulating gene expression in healthy and pathological states of the heart is critical for understanding cardiac biology and disease. By performing enhancer discovery by ChIP-seq on heart tissues from a sizable cohort of individuals, we were able to assess the baseline variation in heart enhancer landscape across the population. Importantly, we found that the activity of cardiac enhancers is well conserved across individuals. This observation, together with the underlying comprehensive maps of heart enhancers generated through this study, provides an important reference point for assessing the impact of environmental factors, genetic variation, and disease-associated molecular processes on cardiac gene regulation.

Similar to the high reproducibility of the enhancer landscape in the healthy heart, our systematic assessment of enhancers in DCM, the most common cardiomyopathy, revealed disease-associated activity changes in thousands of enhancers that were highly reproducible across individuals. This insight implies that pathological disease states of the heart are not associated with a general disruption of coordinated gene expression, but instead show a complex, but well-defined shift in genome-wide transcriptional regulation involving a distinct set of distant-acting enhancers.

Finally, our comparison with data from fetal heart tissue uncovered that, beyond the limited number of examples described to date^13,14^, hundreds of fetal genes and more than a thousand fetal enhancers become consistently reactivated in adult heart disease. Our study identified the genomic locations and activation states of this intriguing subset of disease-associated enhancers, offering an entry point for targeted experimental exploration for hundreds of genes with disease-associated activity changes. Reactivation of individual fetal pathways has also been observed in other, non-cardiac disease processes^15–17^. While limited to anecdotal examples, in conjunction with data from the present study, these reports raise the possibility that widespread reactivation of fetal enhancers is a more generalized disease-associated phenomenon.

Taken together, our study highlights how the etiology of a disease process with major epidemiological relevance is tightly interlinked with, and possibly driven by, reproducible activity changes across thousands of enhancers in the human genome. The identification of substantial overlap between disease-associated and developmentally active cardiac enhancers reinforces the notion that modulation of genes and pathways normally active in the embryonic and fetal heart may impact on disease progression^13^, thus providing inspiration for new therapeutic strategies.

## Methods

### Contact for Reagent and Resource Sharing

All plasmids and reagents generated in this study, as well as archived surplus LacZ-stained embryos for selected enhancers are available from the authors. Please contact Axel Visel.

### Human Subjects

All aspects of this study involving human tissue samples were reviewed and approved by the Human Subjects Committee at Lawrence Berkeley National Laboratory (Protocol Nos. 00023126 and 00022756). Procurement of adult human myocardial tissue was performed under protocols approved by Institutional Review Boards at the University of Pennsylvania (Protocol No. 802781) and the Gift-of-Life Donor Program (Pennsylvania, USA). DCM hearts were procured at the time of orthotopic heart transplantation at the Hospital of University of Pennsylvania. Healthy hearts were obtained at the time of organ donation from cadaveric donors. In all cases, hearts were arrested in situ using ice-cold cardioplegia solution, transported on wet ice, and flash frozen in liquid nitrogen within 4 hours of explantation. All samples were full-thickness biopsies obtained from the free wall of the left ventricle. Clinical summary information for adult DCM and control samples is provided in **Supplementary Table 1.** Fetal and embryonic human heart samples were obtained from the Human Developmental Biology Resource at Newcastle University (hdbr.org), in compliance with applicable state and federal laws and with fully informed consent. Embryonic and fetal samples included: one post conception week 8 (Carnegie stage 22) whole heart sample, three post conception week 10 whole heart samples, and one post conception week 17 left ventricle sample. All adult and fetal samples were shipped on dry ice and stored at −80°C until processed.

### Transgenic mouse assays

All animal work was reviewed and approved by the Lawrence Berkeley National Laboratory Animal Welfare and Research Committee. Transgenic assays were performed in *Mus musculus* FVB strain mice and included both male and female embryos. Sample sizes were selected based on our previous experience of performing transgenic mouse assays for >2,000 enhancer candidates^23,39^. Mouse embryos were only excluded from further analysis if they did not carry the reporter transgene or if they were not at the correct developmental stage. Randomization and experimenter blinding were unnecessary and not performed for transgenic assays, as all resulting embryos were treated with identical experimental conditions and no direct comparisons were made between groups of mice.

### ChIP-seq

Chromatin immuno-precipitations were performed as previously described^40^ with some modifications. Briefly, frozen left ventricle tissue was pulverized with a mortar and pestle, resuspended in PBS, and cross-linked with 1% formaldehyde at room temperature for 10 min. Chromatin was sonicated to obtain fragments with an average size ranging between 100-600 bp. Chromatin was incubated for 2h at 4°C with 5 *μ*g of Active Motif H3K27ac antibody (cat#39133, lot 01613007). Protein A and G Dynabeads (Invitrogen) were then added to this chromatin/antibody mixture for 30 minutes at 4°C. Immuno-complexes were sequentially washed. The protein/DNA complexes were eluted in an SDS buffer (1% SDS, 50 mM Tris pH 8.0, 10 mM EDTA) at 37°C for one hour. Samples were treated with Proteinase K at 37°C and reverse-crosslinked overnight. Finally, the DNA was purified on Zymo ChIP clean and concentrate columns (Zymo Research), and the quality was assessed on the Agilent bioanalyzer. The ChIP-seq libraries were prepared using the Illumina TruSeq library preparation kit followed by pooling of libraries. Libraries were pooled and sequenced via single end 50 bp reads on a HiSeq 2500 (6 libraries per lane) or HiSeq 4000 (8 libraries per lane)(Illumina).

ChIP-seq data was analyzed using the ENCODE Uniform Processing Pipelines (https://www.encodeproject.org/pipelines/) implemented at DNAnexus (https://www.dnanexus.com). ChIP-seq data was analyzed using the ENCODE histone ChIP-seq Unary Control Unreplicated (GRCh38) – pipeline version 1.2 (code available from https://github.com/ENCODE-DCC/chip-seq-pipeline). Briefly, reads were mapped to the human genome (GRCh38) using bwa aln v 0.7.10^41^ with settings -q 5 −1 32 -k 2. Duplicate reads were identified and removed using PICARD (https://broadinstitute.github.io/picard/). Non-duplicate reads were used as input for peak calling using MACS 2.0^42^, with the experiment-matched input DNA used as a control. The average enhancer peak was 6,526 bp long.

### RNA-seq

RNA was isolated from homogenized left ventricle using the TRIzol Reagent (Life Technologies). RNA samples were DNase-treated with the TURBO DNA-free Kit (Life Technologies), and RNA quality was then assessed using a 2100 Bioanalyzer (Agilent) with an RNA 6,000 Nano Kit (Agilent). RNA sequencing libraries were made using the TruSeq Stranded Total RNA with Ribo-Zero Human/Mouse/Rat kit (Illumina) according to manufacturer instructions. RNA-seq libraries were subjected to an additional purification to remove remaining high molecular weight products as follows: sample volume was increased to 100 *μ*l by addition of 1X TE buffer or Illumina Resuspension Buffer and then incubated with 60 *μ*l Agencourt AM-Pure XP beads for 4 min. The beads were pelleted by incubation on a magnet, and the entire supernatant was transferred to a tube containing 50 *μ*l of fresh AMPure XP beads and incubated for 4 min. After pelleting the new beads with a magnet, the supernatant was discarded, the beads washed twice with 80% ethanol and the DNA was eluted in 30 *μ*l Illumina Resuspension buffer. The resulting RNAseq libraries were diluted 10×, and their quality and concentration were assessed using a 2100 Bioanalyzer with the High Sensitivity DNA Kit (Agilent) and a Qubit Fluorometer with the Qubit dsDNA HS Assay Kit (Life Technologies). RNAseq libraries were pooled and sequenced via single end 50 bp reads on a HiSeq 2500 (4 libraries per lane) or HiSeq 4000 (6 libraries per lane)(Illumina).

Like ChIP-seq, RNA-seq data was analyzed using the DNAnexus instance of the ENCODE Uniform Processing Pipeline. RNA-seq data was analyzed using the ENCODE RNA-Seq (Long) Pipeline – 1 (single-end) replicate pipeline (code available from https://github.com/ENCODE-DCC/rna-seq-pipeline). Briefly, reads were mapped to the human genome (GRCh38) using STAR align (V2.12). Genome-wide coverage plots were generated using bam to signals (v2.2.1). Gene expression counts were generated for gencode v24 gene annotations using RSEM (v1.4.1).

### Differential binding analysis

Differential peak analysis between two phenotypes (healthy adult vs DCM adult left ventricle; healthy adult left ventricle vs fetal heart) was performed using DiffBind^26^ version 2.10.0. We identified all autosomal differential peaks at least 1,000 bp from a transcription start site using Gencode^43^ version 27 annotations. Peaks present in at least two individuals were defined using the *dba.count* function in DiffBind, and these peaks were used to generate a list of consensus peaks for comparison with downstream differential analysis. We used the DBA_DESEQ2 method to call differential peaks using the multiple testing corrected FDR ≤ 0.05. Final differentially bound datasets were further filtered to have a two-sided P value < 0.01 and |log_2_(fold change)| ≥ 1. To reduce the dimensionality of the dataset and visualize the samples in two dimensions, the master dataset generated by DiffBind (namely the normalized signal quantifications in all the regions tested across all samples) were log-2 transformed and scaled (z-score by region), and then subject to Principal Component Analysis (PCA) using the prcomp package in R. t-SNE^35^ was run using Seurat^44^ version 2.3.4. TMM-normalized counts were log_2_-converted, imported in a Seurat object, and scaled. PCA was then run using regions showing highly variable signal (mean.function = ExpMean, dispersion.function = LogVMR, x.low.cutoff = 0, x.high.cutoff = 14, y.cutoff = 1, num.bin=100). To determine the correct settings for t-SNE visualization, the optimal number of components was determined by JackStraw, resulting in the top 4 components to be used as input for the t-SNE. This was run using the Seurat internal function and perplexity set to values in the following range: [2, 4, 6, 8, 10, 12]. Figures show t-SNE graphs with perplexity of 10.

### Differential gene expression

Differential gene expression analysis (autosomal genes only) between two phenotypes (healthy adult vs DCM adult left ventricle; healthy adult left ventricle vs fetal heart) was performed using edgeR^45^ version 3.20.9. First, genes whose expression was extremely low in most samples were discarded from further analyses (only those genes showing counts per million mapped reads (CPM) ≥ 1 in three or more samples were retained for further analyses). Next, *estimateCommonDisp* and *estimateTagwiseDisp* (prior.df =10) were run to properly handle over-dispersion at the global and single gene level. Across samples, normalization was then performed using TMM, via the function *calcNormFactors*. Differentially expressed genes were determined using the *exactTest* function, as those showing a false discovery rate (FDR) (Benjamini-Hochberg correction) < 5% and a linear fold-change of at least 2, in either direction. The total number of expressed genes includes all genes present in at least two samples with normalized CPM (counts per million) of at least 1. PCA was performed as described in the previous paragraph, using the log_2_-transformed, TMM-normalized expression values as input. PCA results were robust to the choice of the most variable genes used as input (range tested: 1,000 to 10,000). t-SNE^35^ was run using Seurat^44^ as described in the previous paragraph, with the exception of the parameters used to identify highly variable signal (mean.function = ExpMean, dispersion.function = LogVMR, x.low.cutoff = 0, x.high.cutoff = 20, y.cutoff = 1, num.bin=100). The optimal number of PCs for t-SNE was determined to be 5 (by JackStraw).

### Gene enrichment

To identify gene ontologies enriched in differential regions we ran GREAT^24^ version 3.0.0 using default settings, including the association rule: Basal+extension: 5,000 bp upstream, 1,000 bp downstream, 1,000,000 bp max extension, curated regulatory domains included. We used the Lift Genome Annotations tool in the UCSC genome browser (genome.ucsc.edu) to convert hg38 regions to hg19 regions required for GREAT input. We identified the most significant terms by binomial q value using the GO Biological Process and GO Molecular Function ontologies. To identify gene ontologies represented in differentially expressed genes, we used PANTHER^46^ version 10.0 and ran the PANTHER Overrepresentation Test (release 20181113).

### Transcription factor binding site analysis

We ran HOMER^28^ v. 4.10.3 using *findMotifsGenome.pl* with the following parameters: -size −500,500 -len 6,7,8,9,10,12,14. We used a hypergeometic distribution to estimate the P value of the enrichments. Input regions were split into 1-kbp bins prior to analysis.

### Closest transcription factor start site comparison

We calculated the nearest transcription start site upregulated in DCM for all 4,406 peaks with increased binding in DCM. As controls, we ran 200 iterations sampling 4,406 peaks from the consensus set of 38,883 peaks present in at least two adult samples. We calculated the mean and standard error of these data and plotted the 95% confidence interval for these controls. We also ran the corresponding analysis of the nearest transcription start site downregulated in DCM for all 3,101 peaks with less binding in DCM, and calculated a control set with 200 iterations randomly sampling 3,101 peaks from the same set of consensus peaks. We calculated the mean and standard error of these data and plotted the 95% confidence interval for these controls.

### DCM and fetal peak comparison

As a control, we performed 200 iterations to choose random subsets of all 26,117 fetal peaks and intersected these with the differentially bound DCM vs Health peaks. We used these sets to establish the expected overlap between fetal-biased and DCM-biased peaks.

### *In vivo* transgenic reporter assays

Enhancer candidate regions were cloned into an Hsp68-promoter-LacZ reporter vector39 using Gibson cloning^47^ (New England Biolabs [NEB]). Transgenic mouse embryos were generated by pronuclear injection, and F0 embryos were collected at embryonic day 11.5 and stained for LacZ activity as previously described^25,39^. Only patterns that were observed in at least three different embryos resulting from independent transgenic integration events of the same construct were considered reproducible. The procedures for generating transgenic and engineered mice were reviewed and approved by the Lawrence Berkeley National Laboratory (LBNL) Animal Welfare and Research Committee.

### Quantification and Statistical Analysis

Sample numbers, experimental repeats and statistical tests are indicated in figures and figure legends or **Methods** sections above.

### Data and Software Availability

All ChIP-seq and RNA-seq data are available through GEO: GSE126573. Images of LacZ-stained embryos are available from the VISTA Enhancer Browser (enhancer.lbl.gov), and raw images are available on request from the lead authors.

## ACKNOWLEDGEMENTS

This work was supported by National Institutes of Health (NIH) grants R24HL123879 and R01HG003988 (to A.V. and L.A.P.) and R01HL105993 (to K.B.M and T.P.C). Research was conducted at the E.O. Lawrence Berkeley National Laboratory and performed under Department of Energy Contract DE-AC02-05CH11231, University of California. The human embryonic and fetal material was provided by the Joint MRC / Wellcome (MR/R006237/1) Human Developmental Biology Resource (www.hdbr.org). I.B. was funded through an Imperial College Research Fellowship. This project used the Vincent J. Coates Genomics Sequencing Laboratory at UC Berkeley, which was supported by NIH Instrumentation Grant S10OD018174.

## AUTHOR CONTRIBUTIONS

C.H.S., D.E.D., J.B., A.V., and L.A.P. conceived the project. S.L., T.P.C., M.P.M., and K.B.M. provided fetal or adult heart tissues. C.H.S., S.Y.A., and B.J.M. performed the ChIP-seq and RNA-seq. B.J.M., I.P.-F., C.S.N., T.G., M.K., E.A.L., J.A.A., A.N.H., Q.T.P., S.T., and V.A. carried out transgenic validation. C.H.S., I.B, M.J.B., and Y.F.Y performed computational analyses. C.H.S., A.V., L.A.P. and D.E.D., wrote the manuscript with input from the remaining authors.

## Supplementary Figures

**Supplementary Figure 1:**
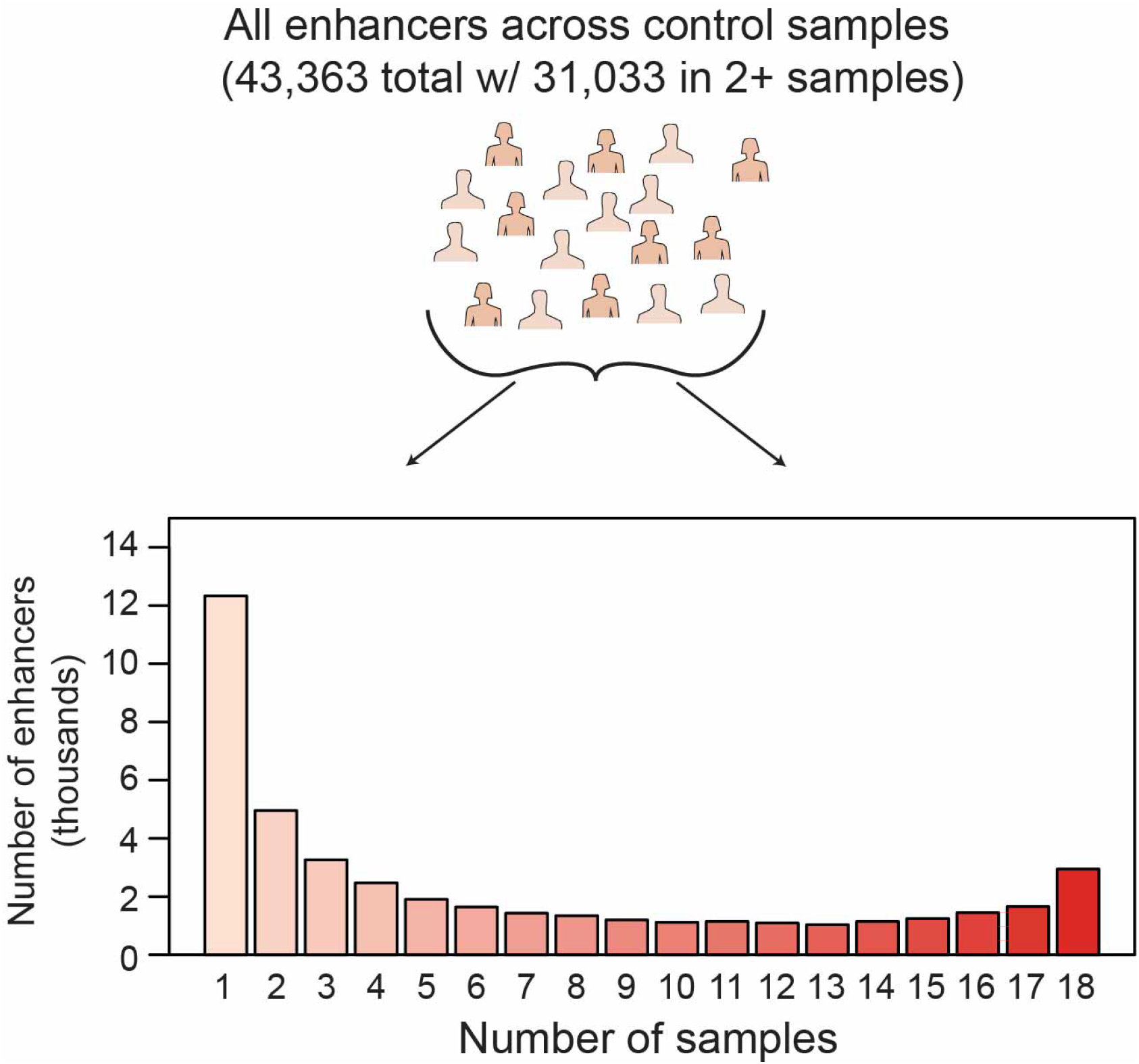
Summary of enhancer representation across all healthy left ventricle samples. Across all healthy samples, we identified 43,363 total predicted enhancers, with more than 31,000 present in at least two subjects

**Supplementary Figure 2:**
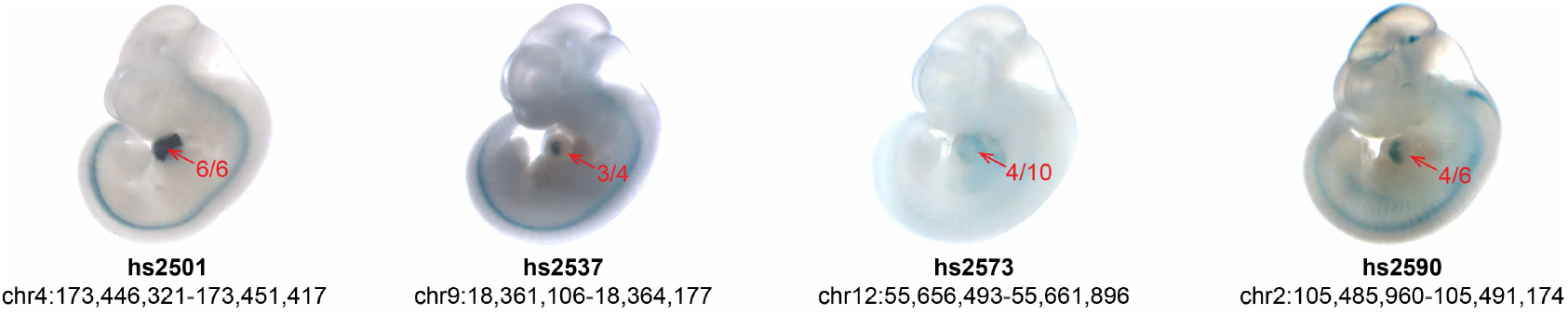
Transgenic validation of additional predicted human heart enhancers. Shown are representative transgenic embryonic day 11.5 (E11.5) mouse embryos showing *lacZ* expression (blue staining) driven by one of four different human heart enhancers. Additional examples are shown in **Figure 2c.** Red numbers indicate reproducibility of the heart expression over the total number of transgene-expressing embryos obtained, hs numbers indicate the unique identifier for the enhancer in the VISTA Enhancer Browser, and the coordinates below are for the tested enhancer in the hg38 assembly of the human genome.

**Supplementary Figure 3:**
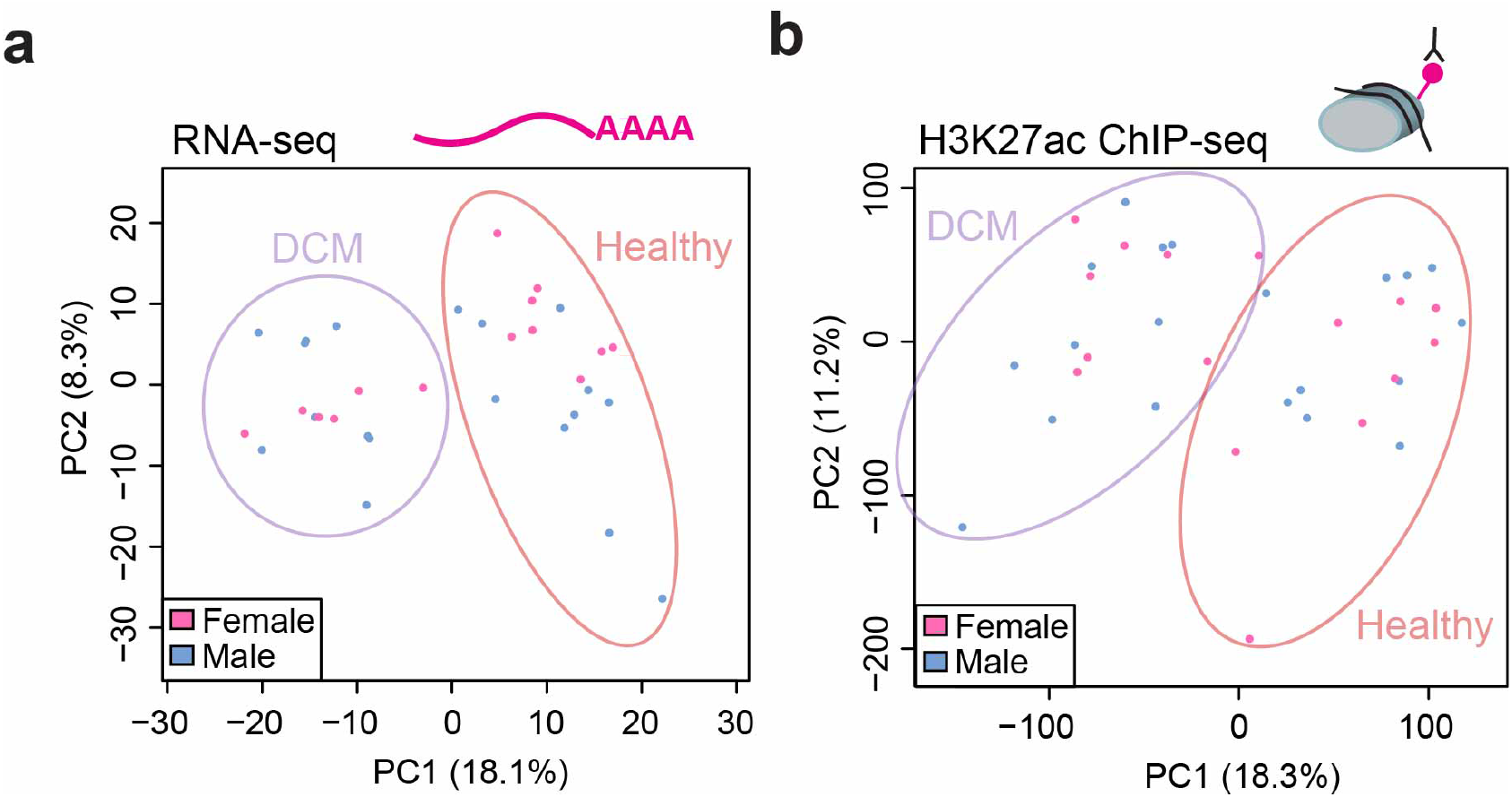
Biological sex is not a major source of variability in heart gene expression or enhancer usage. Principal Components (PC1, x-axes; PC2, y-axes) for expressed genes (a) and distal enhancer peaks (b), as in Figure 3a. In this version, samples from males are colored blue and females pink. These data include only autosomal genes and peaks.

**Supplementary Figure 4:**
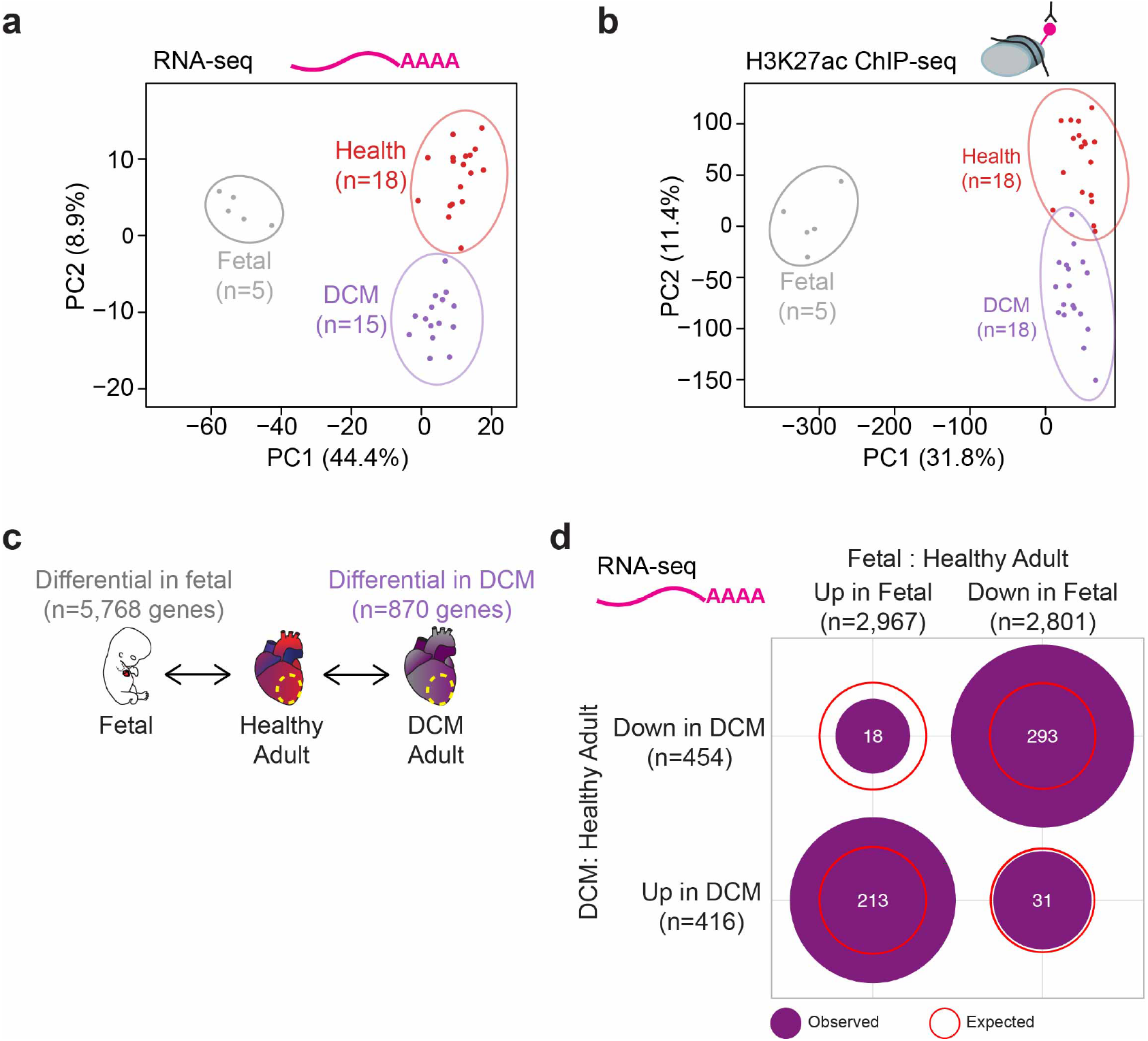
Comparing healthy and DCM adult heart to fetal human heart. **a-b,** Principal Components Analysis (PC1, x-axes; PC2, y-axes) including all adult and fetal human samples for expressed genes (**a**) and H3K27ac enhancers (**b**). Each point represents a unique sample, color-coded by cohort according to legend. PC1 separates developmental stage, while PC2 separates disease state. **c,** Overview of gene expression comparisons: healthy adult versus DCM adult, healthy adult versus fetal. **d,** Number of genes showing correlated or anti-correlated expression between DCM and fetal heart. Filled purple circles indicate observed values, while empty red circles indicate expected values.

**Supplementary Figure 5:**
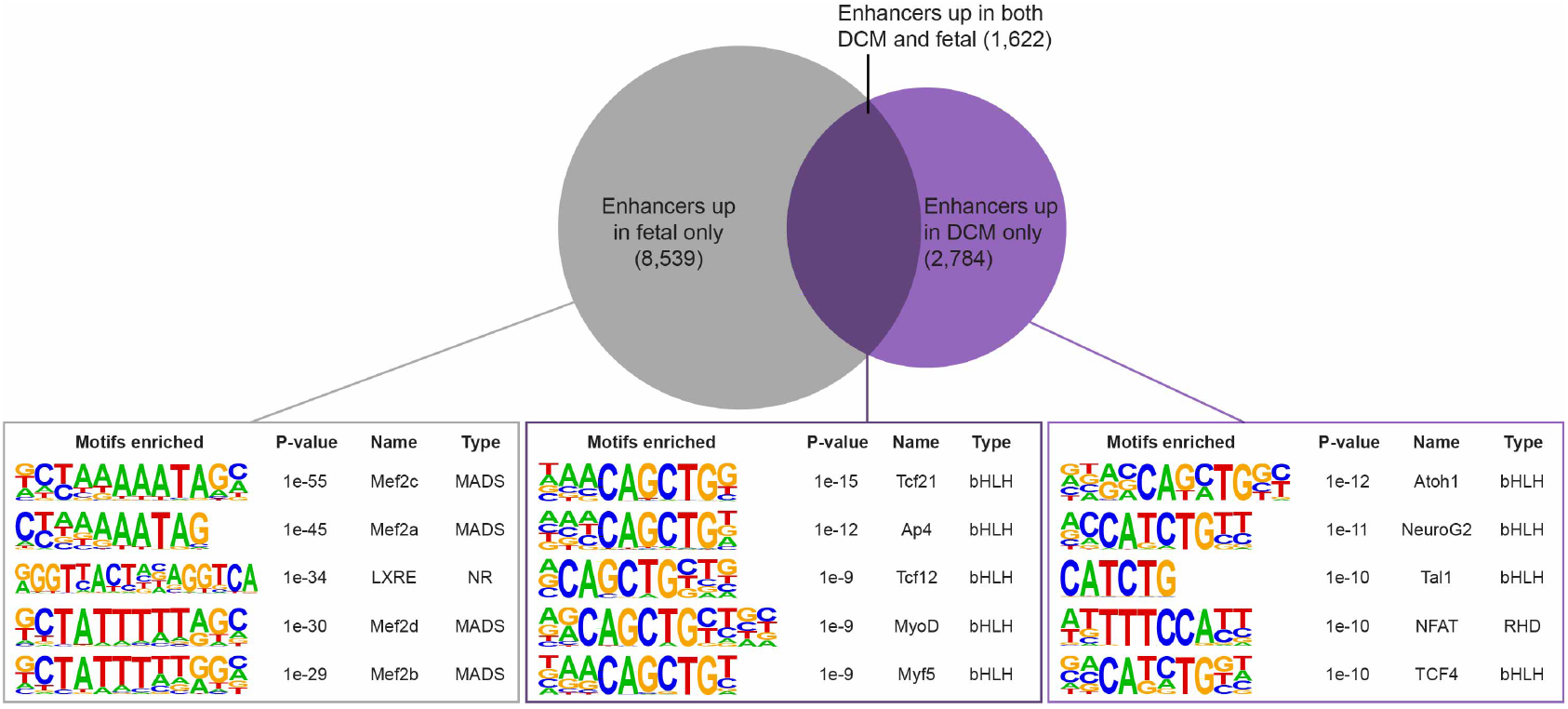
Transcription factor binding motifs enriched in enhancers upregulated in fetal and DCM samples. Shown are the top five most significantly enriched transcription factor binding motifs identified by HOMER (see **Methods**) for enhancers upregulated in fetal samples only (left), enhancers upregulated in both fetal and DCM (middle), and enhancers upregulated in DCM only (right). The background set used for all comparisons was composed of all 12,945 enhancers upregulated in DCM and/or fetal samples. Venn diagram not drawn to scale.

## Supplementary Tables

**Supplementary Table 1.** Clinical characteristics of adult subjects. Quantitative data provided as mean ± standard deviation. * P < 0.05 significantly different between DCM and non-failing cohorts by two-tailed t-test. BMI: body mass index, LV EF: left ventricle ejection fraction.

|  | DCM | Non-failing |
| --- | --- | --- |
| Total subjects | 18 | 18 |
| Gender: Male/Female | 10/8 | 10/8 |
| Age at surgery (years) | 54±8 | 50±10 |
| BMI (kg/m<sup>2</sup>) | 27±6 | 29±7 |
| Prior hypertension (%) | 28 | 33 |
| Prior diabetes (%) | 11 | 5.5 |
| LV EF (%) | 14±3* | 63±6 |
| Heart mass (g) | 525±95* | 373±74 |

**Large tables provided separately as Excel files**

**Supplementary Table 2. Predicted left ventricle enhancers shared by at least two healthy controls.** Predicted enhancers were defined as H3K27ac peaks at least 1 kb away from a transcription start site. A total of 31,033 enhancer peaks were found in at least two controls. Genome coordinates given in hg38.

**Supplementary Table 3. Most highly enriched ontology terms for genes near predicted human heart enhancers.** Analysis was performed using GREAT 3.0 and all peaks present in 16 or more healthy samples. Shown are the 20 most enriched ontology terms from mouse gene expression (MGI expression). Terms in bold are related to cardiovascular tissue.

**Supplementary Table 4. Genes differentially expressed between DCM and healthy heart.** Genes were considered differentially expressed between DCM and healthy hearts with P < 0.01 and |log_2_(fold change)| ≥ 1. In total, 416 genes were upregulated in DCM and 454 downregulated in DCM relative to healthy samples.

**Supplementary Table 5. Predicted left ventricle enhancers shared by at least two adult samples from either cohort.** Predicted enhancers were defined as H3K27ac peaks at least 1 kb away from a transcription start site. A total of 38,883 enhancer peaks were found in at least two adult heart samples (healthy or DCM). Genome coordinates given in hg38.

**Supplementary Table 6. Enhancers differentially bound between DCM and healthy adult heart.** Enhancers were considered differentially bound between DCM and healthy hearts with P < 0.01 and |log_2_(fold change)| ≥ 1. In total, 4,406 enhancers were upregulated in DCM and 3,101 downregulated in DCM relative to healthy samples. Genome coordinates in hg38.

**Supplementary Table 7. Gene ontology terms enriched in differentially active enhancers.** Shown are the top 20 Biological Process terms enriched in enhancer peaks that show increased (top) or decreased (bottom) binding in DCM relative to healthy adult. GO analysis was performed using GREAT.

**Supplementary Table 8. Gene ontology terms enriched in differentially expressed genes.** Shown are the top 20 Biological Process terms enriched in genes that show increased (top) or decreased (bottom) expression in DCM relative to healthy adult. GO analysis was performed using PANTHER.

**Supplementary Table 9. Enhancers differentially bound between fetal and healthy adult heart.** Enhancers were considered differentially bound between fetal and healthy hearts with P < 0.01 and |log_2_(fold change)| ≥ 1. In total, 10,161 enhancers were upregulated in fetal and 15,956 downregulated in fetal relative to healthy adult samples. Genome coordinates in hg38.

**Supplementary Table 10. Genes differentially expressed between fetal and healthy adult heart.** Genes were considered differentially expressed between fetal and healthy hearts with P < 0.01 and |log_2_(fold change)| ≥ 1. In total, 2,967 genes were upregulated in fetal and 2,801 downregulated in fetal relative to healthy adult samples.

**Supplementary Table 11. Gene ontology terms enriched in differentially active enhancers.** Shown are the top 20 Biological Process terms enriched in enhancer peaks that show increased (top) or decreased (bottom) binding in DCM and fetal relative to healthy adult. GO analysis was performed using GREAT.

